# Removal of a genomic duplication by double-nicking CRISPR restores synaptic transmission and behavior in the MyosinVA mutant mouse Flailer

**DOI:** 10.1101/2023.04.28.538685

**Authors:** Fernando J Bustos, Swarna Pandian, Henny Haensgen, Jian-Ping Zhao, Haley Strouf, Matthias Heidenreich, Lukasz Swiech, Benjamin Deverman, Viviana Gradinaru, Feng Zhang, Martha Constantine-Paton

## Abstract

Copy number variations, and particularly duplications of genomic regions, have been strongly associated with various neurodegenerative conditions including autism spectrum disorder (ASD). These genetic variations have been found to have a significant impact on brain development and function, which can lead to the emergence of neurological and behavioral symptoms. Developing strategies to target these genomic duplications has been challenging, as the presence of endogenous copies of the duplicate genes often complicates the editing strategies. Using the ASD and anxiety mouse model Flailer, that contains a duplication working as a dominant negative for MyoVa, we demonstrate the use of DN-CRISPRs to remove a 700bp genomic duplication *in vitro* and *in vivo*. Importantly, DN-CRISPRs have not been used to remove more gene regions <100bp successfully and with high efficiency. We found that editing the *flailer* gene in primary cortical neurons reverts synaptic transport and transmission defects. Moreover, long-term depression (LTD), disrupted in Flailer animals, is recovered after gene edition. Delivery of DN-CRISPRs *in vivo* shows that local delivery to the ventral hippocampus can rescues some of the mutant behaviors, while intracerebroventricular delivery, completely recovers Flailer animal phenotype associated to anxiety and ASD. Our results demonstrate the potential of DN-CRISPR to efficiently (>60% editing *in vivo) remove* large genomic duplications, working as a new gene therapy approach for treating neurodegenerative diseases.

## Background

Genomic recombination events, and particularly copy number variants, have been strongly linked to neurodegenerative diseases such as Alzheimer’s disease and specially autism spectrum disorders (ASD) [1–6]. Targeting these sequences using gene-editing tools has proven challenging as editing a genomic duplication may also affect the endogenous copies of the genes that have been duplicated. Therefore, identifying new strategies for editing genomic duplications without altering the endogenous copies of the wild-type genes holds promise for future therapeutic approaches. In recent years, the discovery and development of CRISPR/Cas9 has enabled versatile manipulation of the genome [7–12]. Among CRISPRs, double-nicking CRISPRs (DN-CRISPRs) have emerged as an alternative to gain on-target specificity in genome editing [13–16]. DN-CRISPRs utilizes SpCas9n (Cas9n), a mutated SpCas9 (D10A; Cas9) that instead of producing double strand breaks, it can only nick one DNA strand, which can be efficiently repaired by the cell without introducing genetic mutations [13]. To successfully edit the genome, it utilizes two sgRNAs that must be located close together on opposite strands of DNA of the desired genomic locus. Since two sgRNAs are needed, this method has the potential for editing genomic duplications without altering the endogenous copies of the wild-type genes. Here, we employ DN-CRISPR to specifically target and remove a genomic duplication in the Flailer mouse model.

The Flailer mouse was identified as a mouse model that carries a spontaneous non-homologous recombination event that resulted in an extra gene, named *flailer,* composed by the fusion of the promoter and exons 1-2 of the *gnb5* gene, fused together by a mixed intron, to exons 26-40 of the *myo5a* gene [17]. The expression of this fusion protein is controlled by the gnb5 gene promoter, which is highly and broadly expressed in the central nervous system (CNS) [18, 19]. The resulting protein, Flailer, binds to cargo but lacks the actin binding domain of MyosinVa, which is necessary for binding and walking to the plus end of actin filaments [20, 21]. In wild-type neurons, MyosinVa plays a crucial role in transporting various components of the synaptic complex, including scaffolding proteins, receptors, and mRNAs, to dendritic spines [22–26]. This process is essential for proper synaptic function and plasticity. When the Flailer protein is present in a 1:1 ratio with wild-type MyosinVa it works as a dominant negative, leading to a reduced transport and abnormal synaptic clustering of receptors and scaffolding proteins such as PSD95 and AMPA receptors [25, 27]. Importantly, removing at least one copy of the Flailer gene could theoretically lead to the recovery of the phenotypes. Previous studies have shown that Flailer neurons exhibit decreased numbers of synaptic spines, increased AMPA receptor-mediated currents, and lack of long-term depression (LTD) in the visual cortex and hippocampus [25, 27]. The enhancement of synaptic activity observed in the FL animal is a result of the absence of LTD. This lack of LTD leads to an inability to prune synapses, resulting in the formation of new synaptic contacts in the dendritic shafts. This is due to the broad distribution of receptors and scaffolds in these regions, as transport to spines is impaired [25]. Furthermore, due to the synaptic defects, Flailer mice display behaviors associated with ASD and anxiety, such as repetitive grooming, seizures, memory deficits, and increased anxiety-like behavior [27]. The Flailer mouse model presents a unique opportunity to dissect the role of different brain regions in controlling behaviors. Since the entire brain of Flailer mice carries the mutation, removing the duplication in specific brain regions allows us to determine the sufficiency of a brain area in controlling a specific behavior and to study its underlying neural circuits.

In this study, we utilized DN-CRISPR technology designing pair of sgRNAs to target exon 2 of *gnb5* and the *myo5a* mixed intron region of the Flailer gene, which encompasses 703bp. By targeting the coding sequence, we aimed to knock out the Flailer gene while also providing the specificity needed to avoid altering the endogenous copies of the genes involved in the duplication. We delivered the sgRNAs via AAV *in vitro* and *in vivo*. Our results showed a reduction in the expression of the Flailer protein and mRNA, and a rescue of the synaptic transmission defects present in Flailer neurons in culture and recovery of LTD in hippocampal slices. Our *in vivo* experiments targeting specifically the ventral hippocampus, show that removal of Flailer allows for partial recovery of anxiety behaviors, while systemic brain delivery of DN-CRISPRs was able to fully recover the behavioral phenotypes exhibited by Flailer animals. These results indicate that the ventral hippocampus alone is not sufficient to fully control anxiety.

Altogether, our findings demonstrate the potential of DN-CRISPRs as a tool for targeting genomic duplications and investigating their effects on neural function and behavior. Moreover, we show that DN-CRISPRs can be used to target large genomic regions with high efficiency, which opens the possibility to new therapeutic strategies.

## Results

### Gene edition of *flailer* gene by DN-CRISPR

As previously described, the *flailer* gene is composed of the first two exons of *gnb5* fused in frame with a mixed intron (*gnb5/myo5a)* and exons 26-40 of *myo5a* (Fig 1A). Since the Flailer mutation is a duplication, every cell also contains the endogenous copies of gnb5 and myo5a [17]. To specifically edit the Flailer gene, we employed DN-CRISPRs, as using CRISPR/Cas9 to edit the mutation/duplication would also affect the endogenous copies gnb5 and *myo5a.* If the endogenous copies of either *gnb5* or *moy5a* are targeted, a new mutation would arise producing a different phenotypes, as such it needs to be avoided. For DN-CRISPR to be able to edit the genome, two sgRNAs need to be close by in opposite strands. This approach proved challenging, as previous studies have demonstrated that the efficiency of editing by DN-CRISPRs is limited, and no evidence of editing was observed when the distance between the two sgRNAs needed to target the genomic locus is greater than 300bp [13, 16, 28].

**Figure 1.**
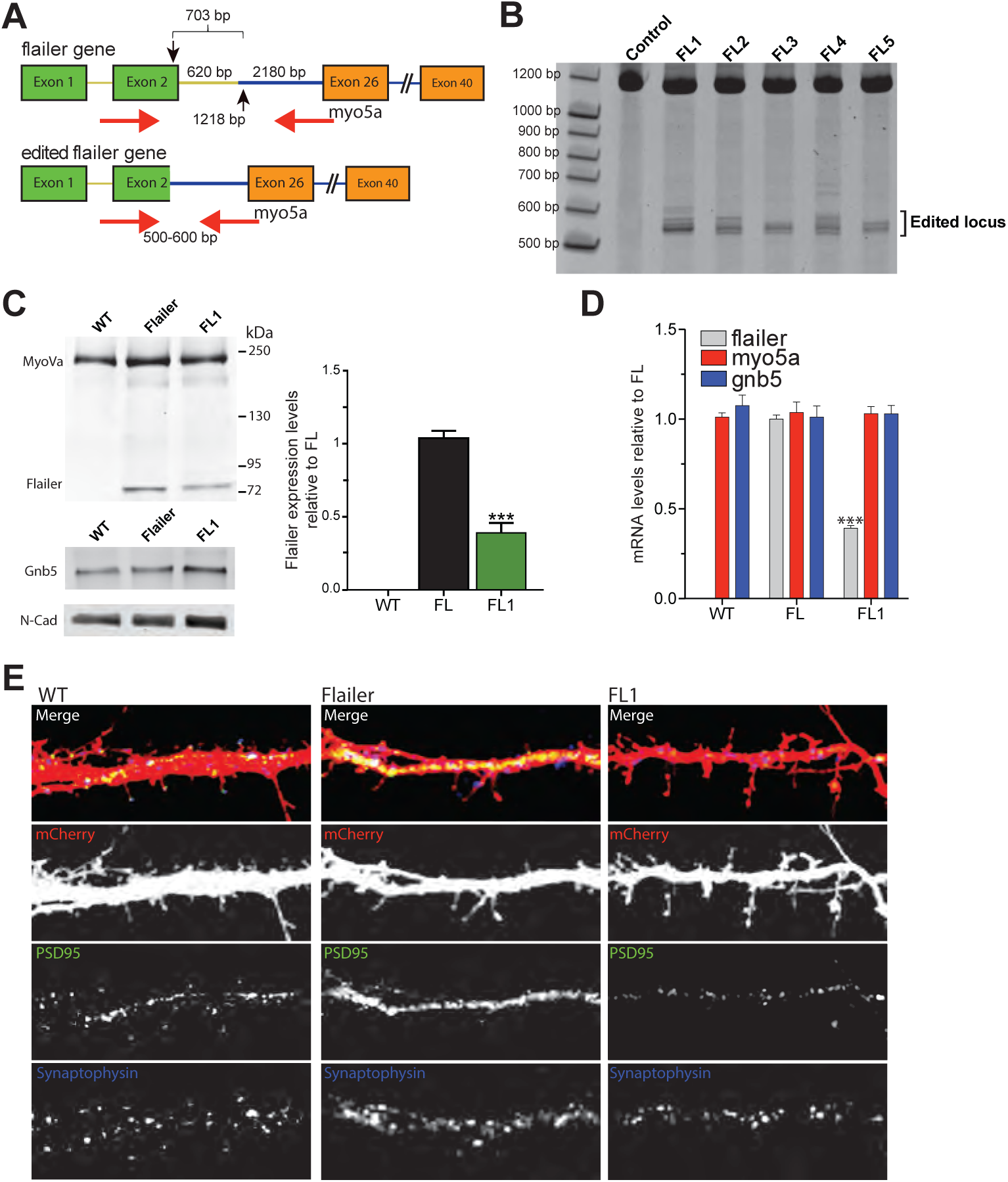
Targeting the Flailer genomic locus with DN-CRISPRs. (A) Schematic representation of the *flailer* gene. Black arrows indicate the region of the DNA targeted by sgRNAs to edit the *flailer* gene via DN-CRISPRs. sgRNAs for DN-CRISPRs need to be targeted at opposite strands. Red arrows show the primers used for PCR analysis to determine the status of the genomic locus. (B) PCR products to analyze gene editing of flailer infected neurons. Protein (C) and mRNA (D) expression levels infected with FL1-FL5. (E) Immunofluorescences against PSD95 and Synaptophysin in wild-type, FL, and FL1-injected neurons. Bars represents mean ±SEM; ***p<0.001. One-way ANOVA was used to determine significance compared to wild-type condition.

Pairs of sgRNAs were designed to target exon 2 of gnb5 and the *myo5a* mixed intron part of the *flailer* gene, a region encompassing approximately 703bp (Fig 1A; black arrows). By targeting the coding sequence we aimed to knock out the *flailer* gene so we can recover the FL phenotype. To identify the most efficient gRNA pair for *flailer* editing, primary cortical neurons were transduced via AAV delivery at 4 DIV with TdTomato alone or in combination with Cas9n and five different sgRNA pairs (FL1-FL5). Six days after infection, genomic DNA was extracted and amplified by PCR using specific primers targeting the *flailer* mutated region (Fig 1A; red arrows). Neurons carrying the complete *flailer* gene are expected to produce a 1218 bp PCR product (Fig 1A; top-red arrows), while edited Flailer gene sequences produce a 500-600 bp PCR product (Fig 1A; bottom-red arrows). As expected, control neurons infected only with Cas9n and TdTomato show a single PCR product of 1218 bp, while neurons co-infected with Cas9n and sgRNAs (FL1-5) using the PHP.eB capsid exhibit a collection of PCR products at ∼500-600bp, indicating successful editing of the locus (Fig 1B). All subsequent experiments were conducted using the FL1 pair of sgRNAs, as it demonstrated the highest efficiency in editing the Flailer genomic locus. PCR amplification products were purified and sequenced, showing the indels produced at the targeted location (Supplementary Fig 1A). In addition, locus-specific sequencing and surveyor assays of the endogenous *gnb5* and *myo5a genomic* sequences that are also nicked by FL1 + Cas9n, revealed that genomic regions were not altered by DN-CRISPR but indel mutations were detected when targeting with wt-SpCas9 (Supplementary Fig 1B-E). To determine the impact of gene editing on FL, we quantified mRNA and protein levels. FL neurons infected for 10 days with FL1 show a robust 70% reduction in mRNA and protein levels compared to FL neurons (Fig 1C-D). As expected, since *gnb5* and *myo5a* genomic locus remained unedited, we did not observe any changes in the mRNA and protein expression levels of wild-type MyoVa or Gnb5 in FL1 treated cells (Fig 1C-D).

We have previously shown that FL cultured neurons exhibit impaired transport of proteins to the synapse, resulting in a lack of the typical clustering structure of receptors and scaffolding protein such as PSD95 and AMPA receptors, hence receptors are distributed extensively in the dendrite shafts where they are able to form functional synapses [25, 27]. To analyze whether editing the *flailer* gene rescues the transport of synaptic proteins and does not show a broad distribution along dendrites of the cargo, we performed immunofluorescence staining using antibodies against Synaptophysin (pre-synaptic protein) and PSD-95 (post-synaptic protein). We found that, similar to wild-type neurons, FL1-treated FL neurons display defined clusters of PSD-95 and Synaptophysin, compared to Flailer neurons, where the expression of PSD-95 is distributed broadly throughout the dendritic shafts and opposing to Synaptophysin signal after 10 days of infection (Fig 1E). Taken together, our data demonstrate that we are able to edit the *flailer* genomic locus, resulting in a reduction of its expression that can rescue the synaptic transport defect in FL cultured neurons.

### Synaptic transmission defects in FL neurons are recovered after *flailer* gene editing

Due to the defects in synaptic transport and the lack of LTD caused by the dominant negative nature of the FL protein, FL neurons produce more synapses localized in the dendritic shafts and in consequence show increased synaptic transmission [25, 27]. To determine whether editing the *flailer* gene can recover synaptic transmission defects, we conducted patch-clamp recordings and calcium imaging of wild-type, FL, and FL1 cultured neurons. Patch-clamp recordings show that the amplitude and frequency of mAMPA receptor currents, enhanced in FL neurons, decrease in FL1 neurons comparable to wild-type cells (Fig 2A-C). Furthermore, current-clamp recording to measure spiking of neurons revealed that FL1-treated neurons have spike frequencies similar to wild-type neurons (Fig 2D-E). As expected, we observe an increase in spiking frequency with development from 4-8 days after infection (Fig 2E). Interestingly, spike frequency was restored to wild-type levels as early as 4 days after infection was performed to correct the mutation (Fig 2E). This finding is supported by calcium imaging using the genetically encoded probe GCaMP6, which shows that FL1-treated neurons have calcium event frequencies that are similar to wild-type neurons and reduced compared to FL neurons as soon as 4 days after infection (Fig 2F). Like current clamp recordings, calcium imaging shows that synaptic activity of neurons increases with development 4-8 days after infection (Fig 2F).

**Figure 2.**
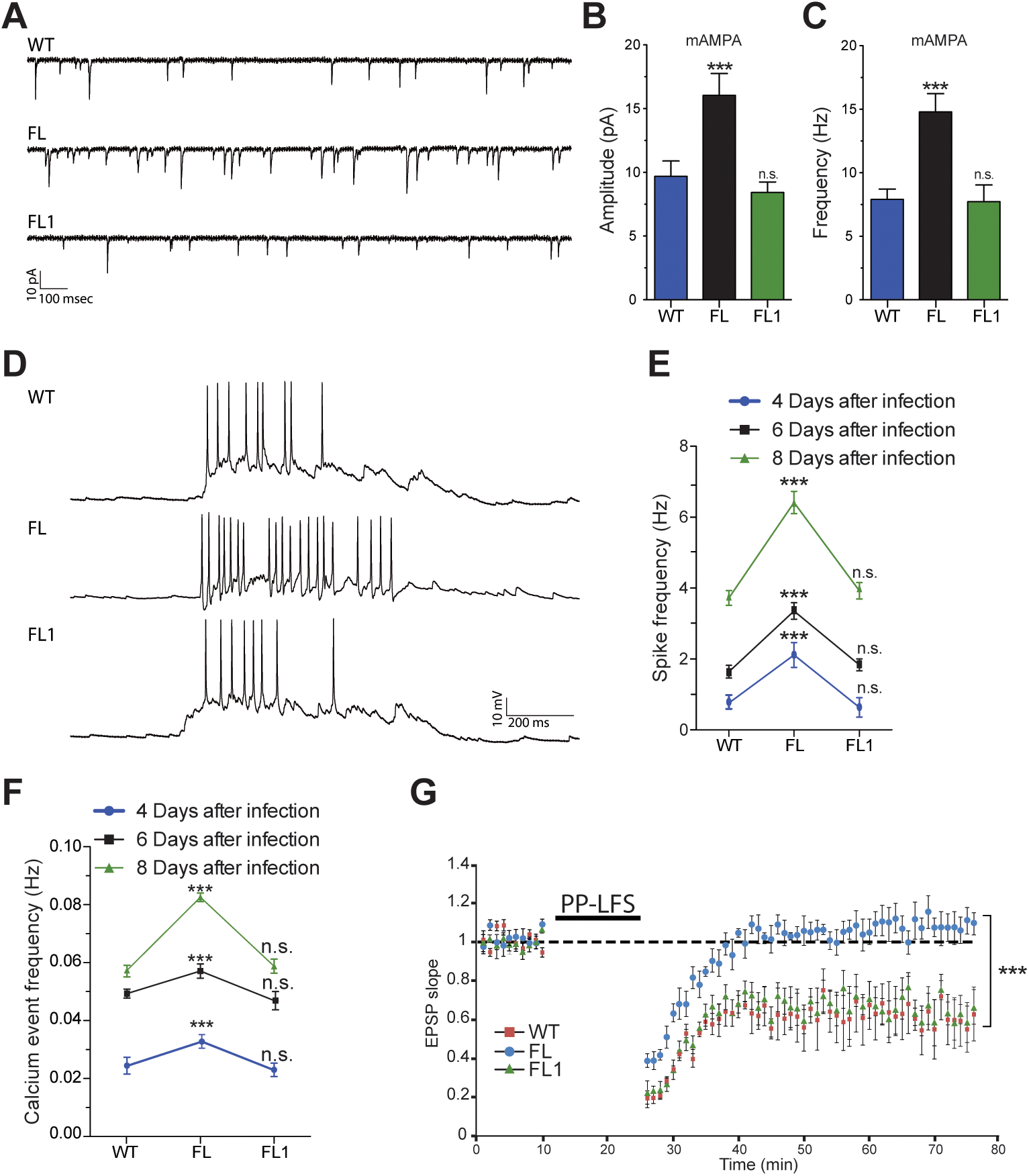
Synaptic transmission defects are corrected by *flailer* gene edition. (A) Representative traces of miniature AMPA receptor currents from wild-type, FL, and FL1 infected neurons. Quantification of the Amplitude (B) and Frequency (C) of currents. (D) Representative traces of current clamp recordings showing spiking of neurons. (E) Spike frequency of wild-type, FL, and FL1 neurons at 4, 6 or 8 days after infection. For A-E At least 20 neurons for each condition from 3 independent neuronal cultures were analyzed. (F) Calcium imaging using GCamp6 at 4-6-8 days after infection. A total of 500 neurons from at least 3 independent neuronal cultures were analyzed. (G) LTD induction by PP-LFS in hippocampal slices of wild-type, FL and FL1 animals at P22-23 after HSV infection. 9 slices from 3 different animals were used for each condition. Bars represents mean ±SEM; ***p<0.001. One-way ANOVA was used to determine significance compared to wild-type condition.

Previously we have shown that FL animals exhibit normal long-term potentiation (LTP), but the induction of long-term depression (LTD) by paired-pulse low-frequency stimulation (PP-LFS) in both the visual cortex [25] and hippocampus [27] is absent. To investigate this further, we used viral delivery to express mruby3 alone or in combination with FL1 and Cas9n into the hippocampus at P20. After 48-72 hours, acute hippocampal slices from wild-type, FL, and FL1 animals were prepared and recorded. As previously reported, FL slices were unable to produce LTD, while FL1 and wild-type slices displayed robust LTD induction by PP-LFS (Fig 2G). Taken together, our data demonstrate that the editing of *flailer* gene with FL1 can restore synaptic function *in vitro* and *in vivo* to levels comparable to those of wild-type neurons.

### Targeted DN-CRISPR editing of the *flailer* gene in the ventral hippocampus ameliorates some of the behavioral phenotypes in FL animals *in vivo*

We have previously characterized the behavioral phenotype of Flailer animals, demonstrating early onset epilepsy, memory deficits, repetitive behaviors, and increased anxiety [27]. Based on our findings *in vitro* and *in vivo* that FL1 corrects synaptic transport and transmission, we aimed to determine if delivering FL1 into a specific brain region could ameliorate some of the behavioral phenotypes associated with the *flailer* mutation, known to be defective. We targeted the ventral hippocampus for gene editing, as this region has been widely implicated in anxiety and memory formation [29, 30]. Tdtomato alone or in conjunction with FL1 and Cas9n was delivered bilaterally via stereotactic injections at P24-P26 (Fig 3A). Four weeks later, animals were tested for behaviors and then sacrificed to determine the accuracy of injection location, percentage of gene edition, and changes in Flailer expression.

**Figure 3.**
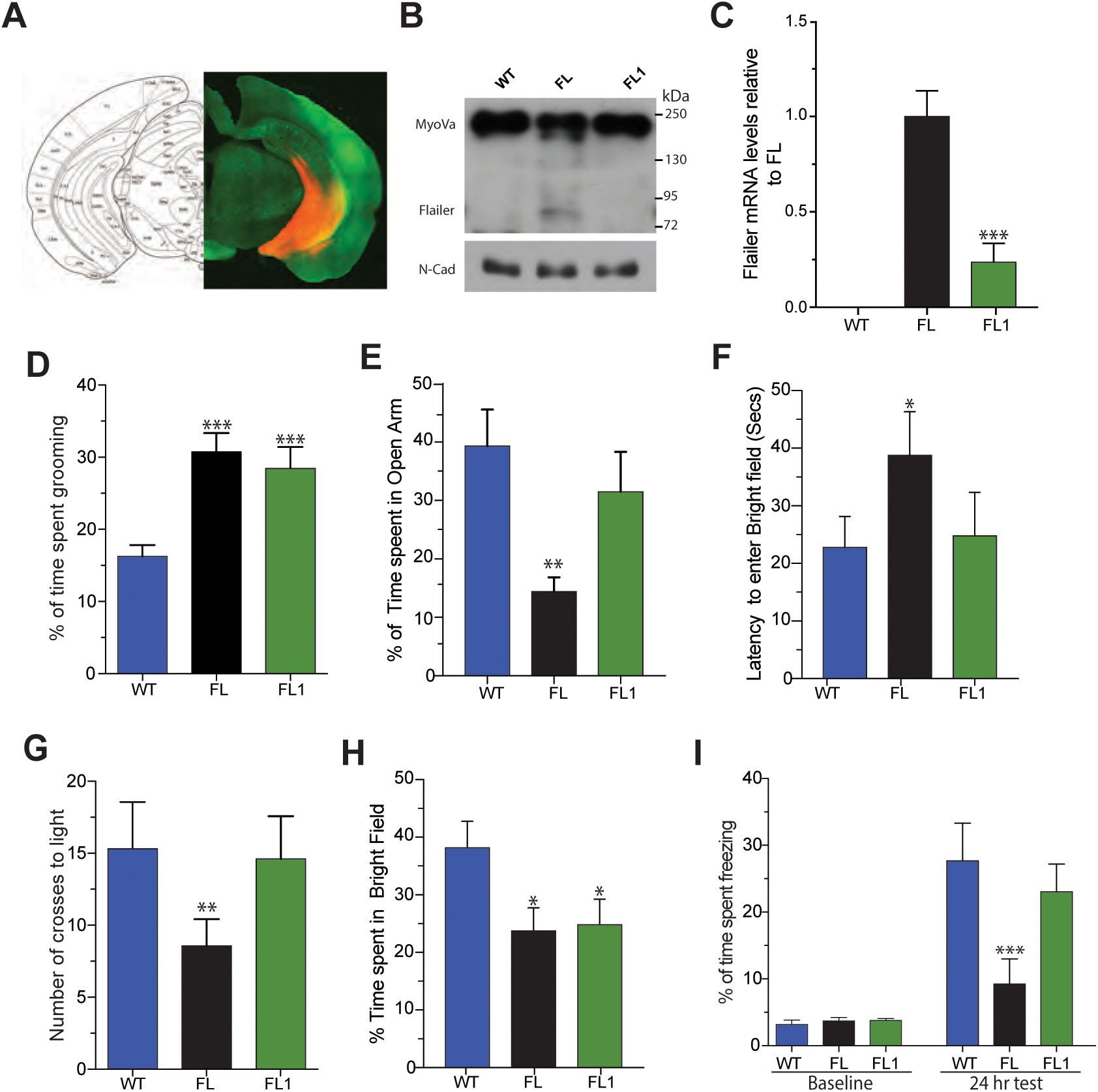
FL1 AAV injection into the ventral hippocampus recovers memory formation and partly anxiety behaviors of FL animals. (A) Representative image of ventral hippocampus stereotactic injection showing high infection rate by TdTomato signal at the targeted region. Infected ventral hippocampus tissue was extracted to determine Protein (B) and mRNA (C) expression levels from wild-type, FL and FL1-injected animals. mRNA levels are relative to FL animals and GAPDH was used to normalize. (D) Percentage of time in a 1 hour session that animals spend grooming. (E) Elevated plus maze test to determine percentage of time spent in the open arms. (F-H) Light and dark test measuring latency to enter the bright field (F), number of crosses to light (G), and percentage of time in bright field (H). (I) Contextual fear conditioning test to measure long term memory formation (24 hours). N=10 animals per condition were analyzed. Bars represents mean ±SEM; *p<0.05, **p<0.01, ***p<0.001. One-way ANOVA was used to determine significance compared to wild-type condition.

To determine changes in Flailer expression, we isolated infected ventral hippocampal tissue from WT, Fl and FL1-injected animals. We found that FL1-injected animals show a 70% reduction in Flailer protein (Fig 3B) and mRNA levels (Fig 3C). To analyze the efficiency of edition, we extracted DNA from infected the ventral hippocampal tissue and run single cell PCRs. A ∼1200bp band indicates the presence of wild type *flailer* gene, while a 500bp indicates that the genomic locus was edited (Supplementary Fig 2A). A total of 500 single cells from 10 FL1-injected animals were used for single-cell PCR showing that 66.3% of the total number of cells analyzed were edited in both copies of *flailer,* 11.3% of neurons only had one copy edited, and 22.4% were not edited at all (Supplementary Fig 2B-C). Only animals that had a clear ventral hippocampal expression of fluorescent proteins and at least a 60% reduction in the expression of Flailer were included in the behavioral analysis.

To analyze the recovery of behaviors, we started investigating grooming behavior. As reported, FL animals groom 30% of their time [27], which is significantly more time than wild-type animals. Animals injected with FL1 in the ventral hippocampus spend a similar percentage of time grooming as FL animals, showing that the condition is not reverted in the FL1-injected animals (Fig 3D). To asses anxiety behaviors, we employed the elevated plus-maze and light and dark tests. The elevated plus-maze test evaluates the time an animal spends in the open or closed arms. If an animal is anxious, it will spend more time in the closed arms. Our data revealed that FL1-injected animals spent more time in the open arms compared to FL animals, at levels comparable to wild-type animals (Fig 3E), showing that FL1-injected animals are less anxious that FL littermates.

The light and dark test consists of two chambers one dark and highly illuminated connected by a door. Anxious animals tend to stay in the dark side and do not come out to explore, thus measuring these parameters can give an estimation of anxiety behavior. We found that FL1-injected animals entered the light side faster and more frequently than FL animals, at levels comparable to wild-type animals (Fig 3F-G). Interestingly, FL1 animals did not stay in the bright field as long as wild-type animals (Fig 3H), indicating that removing the FL mutation in the ventral hippocampus alone is not sufficient to fully recover the anxiety behaviors displayed by Flailer animals. Next, we evaluated long term memory formation (24 hours) using the contextual fear conditioning paradigm. We have previously shown that FL animals are not able to form long term memory in the contextual fear conditioning apparatus, measured by the significant reduction on freezing behavior [27]. We found that similar to wild-type animals, FL1-injected animals spent a significantly more time freezing than FL animals (Fig 3I), which do not form a strong memory as previously described [27]. Together, our results demonstrate that the editing of the *flailer* gene in the ventral hippocampus can significantly improve memory formation and partially alleviate anxiety levels in Flailer animals.

### Intracerebroventricular injection of DN-CRISPRs fully recovers behaviors in FL animals

Given that injection of FL1 in the ventral hippocampus of FL animals was able to improve memory formation and alleviate some anxiety behaviors, we investigated whether whole-brain editing of the *flailer* gene via intracerebroventricular injection could fully recover the characteristic behaviors of Flailer animals [27]. Recently, it has been demonstrated that the delivery of the CRISPR/Cas9 system via AAV packed using the PHP.eB capsid [31], which efficiently crosses the blood-brain barrier and infects the CNS, can ameliorate Alzheimer’s-related phenotypes in an APP Swedish mutation mouse model [32]. We packaged tdTomato alone or together with FL1 and Cas9n (FL1icv) into AAVs using the PHP.eB capsid and delivered them intracerebroventricularly to wild-type and FL animals at P0.

Eight weeks after the injections, behavioral tests were conducted to evaluate anxiety and memory formation. After the tests, the animals were sacrificed to assess infection efficiency, Flailer expression, and editing efficiency. As expected, infection with AAV using the PHP.eB capsid produced a strong infection shown by TdTomato signal, concentrating more in the hippocampus and cortex, but also widely distributed throughout the brain (Fig 4A-B). Using infected cortical tissue, we determined the expression levels of Flailer after edition. FLicv animals exhibited a significant 70% reduction in the expression of the Flailer protein (Fig 4C) and mRNA levels (Fig 4D), compared to FL littermates. To analyze the edition efficiency produced in infected cortical tissue, single-cell PCR from 500 cells obtained from FL1icv injected animals showed that 61.7% of the total number of cells analyzed were edited in both copies of *flailer,* 10.6% of neurons only had one copy edited, and 27.6% were not edited at all (Supplementary Fig 2D-E). Only animals that had high and broad infection and at least a 60% reduction in the expression of Flailer were included in the behavioral analysis.

**Figure 4.**
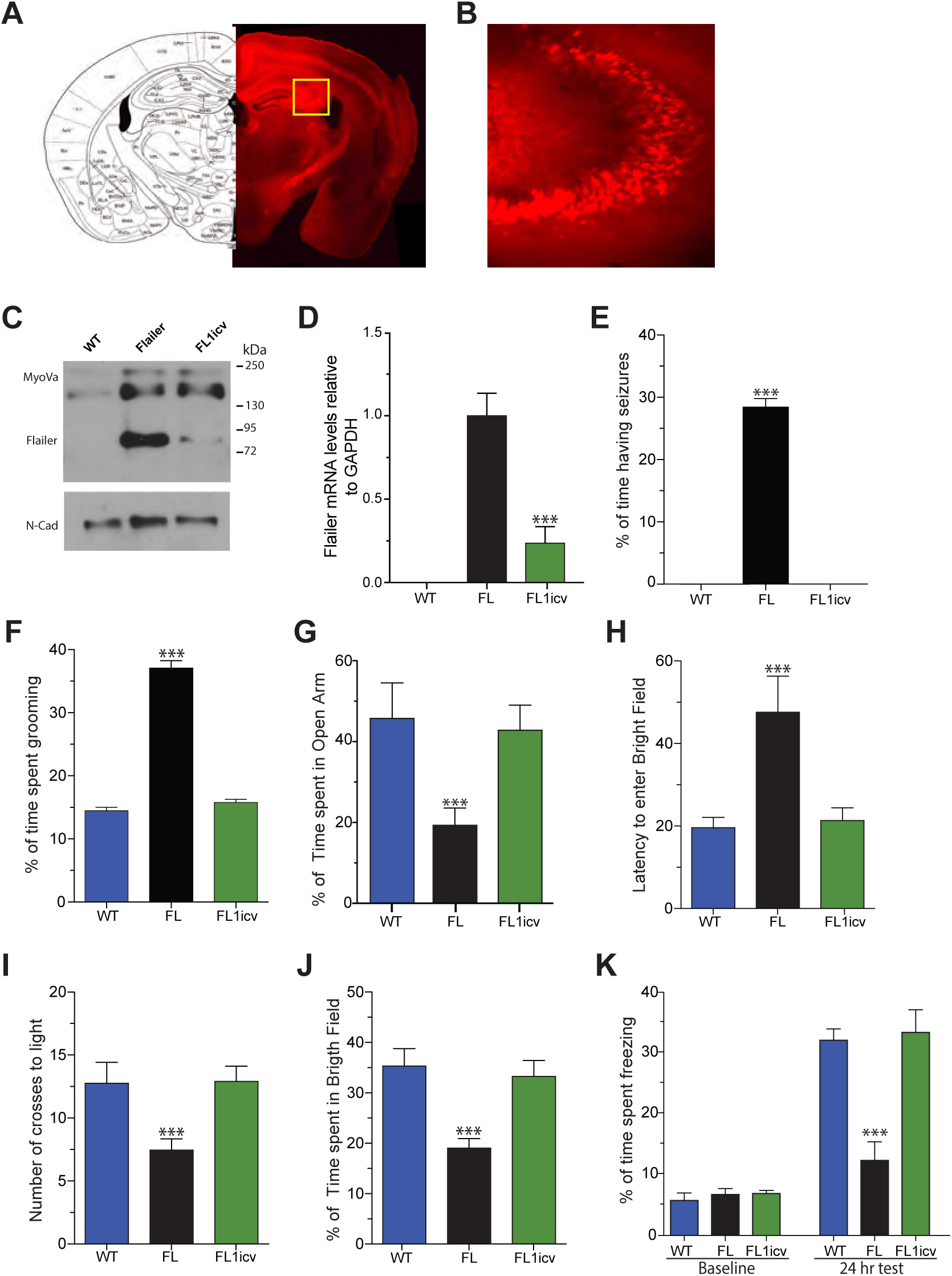
Intracerebroventrical injection of FL1 recovers ASD and anxiety-like behaviors of FL mice. (A) Representative image of intracerebroventricular injection showing high and extensively distributed infection by TdTomato signal. (B) Magnification of the hippocampus area depicted in a yellow square in A. Protein (C) and mRNA (D) expression levels of Infected tissue extracted from the cortex of wild-type, FL and FL1icv-injected animals. (E) Quantification of the time animal spend having seizures. (F) Percentage of time in a 1 hour session that animals spend grooming. (G) Elevated plus maze test to determine percentage of time spent in the open arms. (H-J) Light and dark test measuring latency to enter the bright field (H), number of crosses to light (I), and percentage of time in bright field (J). (K) Contextual fear conditioning test to measure long term memory formation at 24 hours. N=10 animals per condition were analyzed. Bars represents mean ±SEM; ***p<0.001. One-way ANOVA was used to determine significance compared to wild-type condition.

Flailer animals are known to exhibit severe seizure episodes until around P30, at which point they suddenly stop [27]. To determine the percentage of time animals have seizures, we recorded them for 1 hour and manually calculated the time spend seizing at P20-P25. We observed that FL1icv animals showed no signs of seizures at P20-P25, in contrast to Flailer littermates who displayed seizures for approximately 30% of the time analyzed (Fig 4E). Next, we analyzed grooming behavior by putting the animals in a cage for 1 hour and recording their behavior. As we have previously shown, FL animals spend 35% of their time grooming [27], in contrast to wild-type and FL1v animals, which groomed approximately 15% of their time, indicating a reversal of the condition (Fig 4F).

To analyze the presence of anxiety behaviors, we used both the elevated plus maze and light and dark tests. We have shown that in both these tests, FL animals have increased anxiety compared to wild-type animals [27] In the elevated plus maze test, we found that FL1icv animals spent more time in the open arm compared to FL littermates and at levels comparable to wild-type animals (Fig 4G). Using the light and dark test, we found that FL1icv animals had a reduced latency to enter the bright field compared to FL animals and at levels comparable to wild-type animals (Fig 4H). In addition, FL1icv animals, similarly to wild-type animals, showed a significant increase in the number of crosses (Fig 4I) and the percentage of time spent in the light compartment (Fig 4J), compared to FL littermates. Lastly, we tested the animals for long-term memory formation at 24 hours using the contextual fear conditioning paradigm. We found that FL animals showed reduced freezing times compared to wild-type animals, while both wild-type and FL1v animals showed comparable levels of freezing (Fig 4K). Our results demonstrate that intracerebroventricular injection of AAVs carrying the FL1 gene editing construct using the PHP.eB capsid can effectively and efficiently recover the characteristic behaviors of Flailer animals, including the reduction of seizures, repetitive behaviors, memory deficits, as well as alleviating anxiety behaviors.

## Discussion

Many neurodegenerative diseases, including ASD and Alzheimer’s disease, have been strongly linked to genomic rearrangements, including gene duplications that result in abnormal gene dosage [1, 2, 33–35]. Even though the use of CRISPR/Cas9 has proven successful targeting different genes in mice to remove a specific genomic sequence [32, 36–38], developing gene therapy approaches to target genomic duplications has proven challenging as a high degree of specificity is needed to avoid targeting the endogenous copies of the duplicated genomic sequence. Moreover, some duplications have structures where large fragments of DNA need to be removed in order to silence gene expression. Here, we employ the CRISPR/Cas9 system, specifically DN-CRISPRs [13–16], to precisely target a genomic duplication that produces ASD and anxiety-like behaviors in mice. One limitation of using DN-CRISPRs for gene editing is the requirement for sgRNAs to be in close proximity. Previous studies have shown that when sgRNAs are more than 100bp [13, 39] or at most 300bp [16] apart from each other, excision efficiency is significantly reduced. Here we demonstrate the feasibility of using DN-CRISPR to target and remove a large (>700bp) genomic duplication, specifically the Flailer duplication that results in a reduction of Flailer expression *in vitro* and *in vivo.* Importantly, we show that our approach is specific as the endogenous copies of *gnb5 and moy5a* that are in part targeted by FL1 DN-CRISPRs, were not edited and no changes in their expression were observed. Our results demonstrate for the first time that increasing the offset between sgRNAs in DN-CRISPRs can enable the precise targeting and removal of large genomic regions, expanding the potential for gene editing to treat genomic duplications associated with neurodegenerative diseases such as ASD, where gene duplications in genes like SHANK3, NRXN1, or NLGN1 have been implicated [1– 6]. This approach could potentially be applied to other genetic disorders characterized by genomic duplications, offering a new avenue for the development of gene therapies.

The Flailer animal was first described by Jones et al., (2000) showing its role as a dominant negative of MyoVa. In wild-type neurons MyoVa moves to the positive end of actin filaments [20], transporting key components to the synapse such as receptors, scaffolding proteins, and mRNAs [23–26]. The Flailer mutation impairs the ability of MyoVa to transport these components, resulting in decreased numbers of synaptic spines, increased AMPA receptor-mediated currents, and the lack of LTD [25]. Our experiments in vitro demonstrated that after editing the Flailer gene, the capacity of transport of PSD95, a key scaffolding protein in the synapse, were recovered observing clusters of PSD95 similar to those in wild-type neurons.

Our experiments have revealed that the Flailer animal model exhibits increased synaptic activity and reduced LTD [25, 27], which is consistent with other ASD models [40–42]. This suggests that the failure of LTD to properly eliminate synapses may contribute to this phenotype. By targeting the Flailer gene using DN-CRISPRs, LTD induction was recovered in hippocampal slices *in vivo* and a recovery of spontaneous activity and spiking of neurons was observed (Fig 2A-F). The failure of LTD in Flailer and other mutant mouse models of TSC1, Pcdh10, or Pten has been consistently linked to the development of ASD [27, 40, 42, 43]. This dysfunction in LTD leads to a lack of activity-dependent synapse elimination, or “pruning,” during development. The absence of LTD and the resulting hyperconnectivity between neurons suggest that the process of LTD and synapse elimination are particularly susceptible to disruption during brain development.

The use of whole-genome knockouts or conditional knockouts with specific promoters has been a common approach to investigate the role of specific brain regions or cell types in controlling synaptic transmission or behavior. Having the whole brain intact and selectively disrupting the targeted area allows us to test the necessity of a specific brain area in a particular brain circuitry in controlling a particular behavior. Conversely, the Flailer animal model presents a unique opportunity to study the sufficiency of brain regions in controlling specific behaviors. The Flailer protein is expressed broadly in the brain, having a dominant negative effect on MyoVa, resulting in the global disruption of brain function. By using region specific AAV delivery of DN-CRISPR-mediated gene editing, we can selectively recover specific brain regions and investigate if these regions are sufficient to control specific behaviors, as shown for the ventral hippocampus. Through this approach, we can use the Flailer model to identify the neural circuits and brain regions that are involved in behaviors related to human neurological diseases.

Flailer animals exhibit a range of behavioral phenotypes, including early onset epilepsy, anxiety, repetitive behaviors, and memory deficits [27], which resemble those seen in ASD and anxiety-like animal models [44]. We first focused on the ventral hippocampus, a brain region that has been implicated in the regulation of anxiety and memory formation [26,27]. Our experiments targeting the ventral hippocampus showed that memory formation is recovered but the anxiety behavior is only partially restored. This suggests that anxiety behaviors are influenced in part by the ventral hippocampus, but edition of this region alone is not sufficient to fully recover these behaviors through gene editing of the Flailer duplication.

To evaluate the possibility to fully recover the animal phenotype, we turned to intracerebroventricular delivery of FL1 using the high infectivity PHP.eB capsid [31]. This capsid has been used to deliver CRISPR/Cas9 in an Alzheimer’s model, resulting in improvement in cognitive performance and reductions in Alzheimer’s markers such as amyloid-beta deposition [32]. Our results demonstrate that after brain injection of FL1 in Flailer animals, we are able to fully recover all of the tested behaviors, including epilepsy, grooming behavior, anxiety, and memory formation. The full recovery was likely due to the high efficiency of genome editing in vivo, with approximately 70% of at least one copy of *flailer* being edited, an no alteration of the endogenous copies of *gnb5* or *moy5a*. This level of editing is significantly higher than in other studies where DN-CRISPRs [13] or CRISPRs [32] have been used to edit the genome, with an efficiency of approximately 30%. These results demonstrate that gene therapy in mice using DN-CRISPRs is able to efficiently edit the genome, resulting in the recovery of synaptic and behavioral phenotypes in the animals. Remarkably, we were able to reach reversion on most all the phenotypes tested including seizures, grooming behaviors and cognitive tests. This shows the prefeasibility of using DN-CRISPR as an approach for developing therapies of neurological diseases.

Altogether, our findings show that the use of DN-CRISPRs is an effective and efficient method for targeting genomic duplications, thereby opening new opportunities for therapeutic interventions were genomic aberrations need to be amended.

## Conclusions

We expand the use of DN-CRISPRs by showing the removal of a genomic region that is greater than >700bp long. This opens new possibilities for gene editing methodologies combined with the increase specificity conferred by DN-CRISPR. Using AAV viral delivery we demonstrate efficient (>60%) gene edition to remove the Flailer duplication. High levels of edition permit the recovery of ASD/anxiety phenotypes in Flailer mice. Together we show the use of DN-CRISPR as a new therapeutil approach for genomic aberrations.

## Materials and Methods

### Animals

All protocols involving rodents were carried out according to NIH and ARRIVE guidelines. Protocols were approved by MIT Committee on Animal Care (CAC), and Ethical and Bio-security Committees of Universidad Andres Bello. Flailer (Jackson Laboratories Strain #:000502) and wild-type (Jackson Laboratories Strain #:000664) animals were used for behavioral and *in vitro* analyses.

### DNA Constructs

To express Cas9n, the coding sequence of SpCas9n (Streptococcus pyogenes Cas9) was obtained from pX335 (Addgene #42335) and subcloned into a AAV backbone under the MeCP2 promoter as in [45]. 20nt target sequences were selected contiguous to a 5’-NGG photospacer-adjacent motif (PAM) sequence in both sense and antisense strands. sgRNA were targeted to the myo5a mixed intron (intron 3) or the second gnb5 exon in the *flailer* gene (Fig 1A). Each pair of sgRNAs (1 sense/1 antisense strand) were synthesized by gBlocks (IDT, USA) under the human U6 promoter and subcloned into an AAV backbone vector coding for tdTomato or Azurite under the hSyn1 promoter. Both sgRNAs were delivered in the same vector. sgRNA pair sequences were design with the first sgRNA targeting exon 2 of *gnb5* and the second targeting the intron part of *myo5a* for specificity. Sequences are as follows: FL1 (*gnb5*: 5’<colcnt=9> CTTTGCACCAATCCATGCAC 3’; *myo5a:* 5’ ATGTTCATGCTTCTATTGAC 3’); FL2 (*gnb5*:5’ ACAAAGTCCTGTGCATGGAT 3’; *myo5a:* 5’ ATAAAAATTAGTACATGTAT 3’); FL3 (*gnb5*: 5’ TGCACAGGACTTTGTTCCCG 3’; *myo5a:* 5’ ATGTTCATGCTTCTATTGAC 3’); FL4 (*gnb5*: 5’ ACAAAGTCCTGTGCATGGAT 3’; *myo5a:* 5’ CTTACCACGTATAAGATGCT 3’); FL5 (*gnb5*: 5’ CTTTGCACCAATCCATGCAC 3’; *myo5a:* 5’ TGAACCAAGCATCTTATACG 3’). For calcium imaging, a plasmid coding for GCamP6s under the CamKIIa promoter in a AAV backbone was obtained from Addgene (Cat #51086).

### Cell line cultures and transfections

Neuro-2a (N2a) and HEK293T cells were grown in DMEM containing 10% Fetal bovine serum (FBS). Cells were maintained at 37 °C in 5% CO2 atmosphere. Cells were transfected using Lipofectamine 3000 or Polyethylenimine (PEI) “MAX” reagent (Polysciences, Cat 24765), according to manufacturer’s protocols.

### Primary neuron cultures

Postnatal day 0 Flailer or wild-type animals were euthanized by decapitation and the whole brain was extracted in ice cold Ca^2+^/Mg^2+^-free Hank’s balanced salt solution (HBSS) solution. Meninges were removed, the tissue was minced and incubated with Papain (20 units) for 15 minutes at 37°C. Cells were rinsed twice with HBSS, resuspended by mechanical agitation through fire-polished glass Pasteur pipettes of decreasing diameters, and plated over poly-L-lysine-coated culture plates or cover slips. Cultures were maintained at 37 °C in 5% CO2 in growth media (Neurobasal-A (Life technologies 1088802) supplemented with B27 (Life technologies 17504044), 2 mM L-glutamine (Life technologies 25030-081), 100 U/ml penicillin/streptomycin (Life technologies 15070-063)]. Half of the media was replaced every 3 days.

### AAV Production

For infection of cultured neurons, low titer viral particles were produced using AAV1/AAV2 or PHP.eB [31] capsids. Briefly, HEK293T cells were grown to approximately 6×10^4^ cells/cm^2^ with DMEM 10% FBS. Cultures were transfected using PEI “MAX” reagent (Polysciences, Cat 24765) with plasmids coding for the capsids, the transfer vectors (sgRNAs with fluorophores, SpCas9n, or GCamP6), and the helper plasmid pDF6. 24 hours after transfection, the media was exchanged for DMEM 1% FBS. After 72h, media and cells were collected from the plates, centrifuged and cell pellet was lysed by freeze thaw cycles. Supernatant was filtered with 0.45μm syringe filters are stored at −80°C for posterior use. High titer viral particles for injections were prepared as in [46]. Briefly, HEK 293T cells were grown to approximately 6×10^4^ cells/cm2 with DMEM 10% FBS. Cultures were transfected using PEI “MAX” reagent (Polysciences, Cat 24765) with PHP.eB capsid plasmids, the transfer vectors (sgRNAs with fluorophores, or SpCas9n), and the helper plasmid DF6. After 24 hours of transfection, the media was exchanged for DMEM 1% FBS. After 72 hours, media was collected from the plates and replaced with fresh DMEM 1% FBS. The collected media was stored at 4°C. 120 hours after transfection, the cells were detached from the plate and transferred to 250 mL conical tubes, together with the collected media. Cells were centrifuged for 10 min at 2000 g, and the supernatant was removed and saved for later use. The pellet was resuspended in SAN digestion buffer (5 mL of 40 mM Tris, 500 mM NaCl and 2 mM MgCl2 pH 8.0) containing 100 U/mL of Salt Active Nuclease (Arcticzymes, USA) and incubated at 37°C for 1 hour. The supernatant was precipitated using 8% PEG 8000 and 500mM NaCl. It was incubated on ice for 2 hours and centrifuged at 4000 g for 30 min in 250 mL bottles. The supernatant was collected and resuspended with SAN digestion buffer. The solution was placed in an iodixanol gradient and ultracentrifuged at 350,000g for 2.5 hours. The phase containing the AAV was rescued and frozen at −80°C for later use.

### Immunofluorescent staining

Immunocytochemistry was performed as described [25, 47, 48]. Briefly, 10-12 DIV neurons were fixed with 4% paraformaldehyde (PFA) and permeabilized with 0.1% Triton X-100 for 5 min. Cells were blocked with 1% BSA for 30 minutes at 37C and incubated with primary antibodies overnight at 4°C. Primary antibodies used were: PSD95 (Neuromab 75-028), Synaptophysin (Zymed 18-0130), and mCherry (Biorbyt orb153320). Cells were rinsed three times with PBS and incubated for 2–3 h at room temperature with appropriate secondary antibodies conjugated to Alexa-488, Alexa-546 or Alexa-647 (Life technologies). Cells were coverslipped using Prolong Gold and imaged on a Nikon C2+ confocal microscope with a 60x oil immersion objective (Nikon Instruments Inc., Melville, NY, USA).

### Purification of cell nuclei and sorting

Animals injected with AAV coding for FL1 were sacrificed and infected brain tissue was quickly extracted in ice cold DPBS using a fluorescent dissecting scope. Samples were directly used or shock frozen on liquid nitrogen. Nuclei purification was performed as described in [45]. Briefly, dissected tissue was homogenized in 2 ml ice-cold homogenization buffer (HB) (320 mM sucrose, 5 mM CaCl, 3 mM Mg(Ac)2, 10 mM Tris pH7.8, 0.1 mM EDTA, 0.1% NP40, 0.1 mM PMSF, 1 mM β-mercaptoethanol) using 2 ml Dounce homogenize; 25 times with pestle A, followed by 25 times with pestle B. HB was added to complete 5 ml and kept on ice for 5 minutes. 5 ml of 50% OptiPrep density gradient medium containing 5 mM CaCl, 3 mM Mg(Ac)2, 10 mM Tris pH 7.8, 0.1 mM PMSF, 1 mM β-mercaptoethanol was added and mixed. The resulting lysate was loaded on top of 10 ml 29% iso-osmolar OptiPrep solution in a conical 30 ml centrifuge tube (Beckman Coulter, SW28 rotor). Samples were centrifuged at 10,100 × g (7,500 r.p.m.) for 30 min at 4 °C. The supernatant was removed, and the nuclei pellet was gently resuspended in 65 mM β-glycerophosphate (pH 7.0), 2 mM MgCl2, 25 mM KCl, 340 mM sucrose and 5% glycerol. Extracts were controlled for purity using bright field microscopy. Purified intact nuclei were sorted into single-cell nuclei using BD FACSAria III (Koch Institute Flow Cytometry Core, MIT). Nuclei were sorted in 5 μl of QuickExtract DNA Extraction Kit (Epicentre, USA) in 96-well format.

### Single cell PCR

Extracted DNA from sorted single nuclei from Flr and Flr-FL1 neurons was used for PCR analyses to determine edition of alleles in single cells. PCR was conducted using nested PCR primers with a first amplification step of 40 cycles using primers: 5’ AAGAAAGCTTTGCATGCCCT 3’, 5’ AATGGCTCCAAACCAACACA 3’; followed by 25 cycles with a second set of primers: 5’ AGCTTTGCATGCCCTAAGAA 3’, 5’ CTCCAGGGCAATTGTAGCCA 3’. Samples were run in agarose gels to visualize allele edition (Supplementary Fig 2).

### Genomic DNA extraction and edition efficiency

Primary cortical neurons were infected at 3 DIV with FL1 and at 10-12 DIV neurons were harvested and DNA extracted using Quick-DNA Miniprep Kit (Zymo Research, USA) following manufacturer recommendations. For tissue, injected animals with AAV coding for FL1 were sacrificed and infected brain tissue was quickly extracted in ice cold PBS using a fluorescent dissecting scope. Tissue was digested with Proteinase K and purified using Quick-DNA Miniprep Kit (Zymo Research, USA) following manufacturers protocol. To test Flailer edition PCR with primers encompassing the edited region (5’ AGCTTTGCATGCCCTAAGAA 3’, 5’ CTCCAGGGCAATTGTAGCCA 3’) were used to determine edition of the locus. PCR products were run in agarose gels and purified by Gel extraction kit (Qiagen). DNA was purified cloned into a TOPO-TA (Lifetechnologies, USA) and sent for Sanger sequencing to corroborate edition of the gene. To test specificity of sgRNA against Flailer, SURVEYOR assay against *gnb5* or *myo5a* was performed using SURVEYOR nuclease assay (IDT, USA) using the following primers: *gnb5* sense 5’ AATTGTGGTCCTGTCCTTGC 3’, an|sense 5’ TGGGTTCCTTCACAAATCCT 3’, myo5a sense 5’ TTACAATGAAGACGCTGTGGA 3’, antisense 5’ CTTTACTGTCCCCACCATGG 3’.

### Electrophysiology

Whole cell patch-clamp recordings were performed on Flr and wild-type-Flr primary cortical neurons as previously described [25, 49, 50]. Briefly, neurons were transferred to external solution containing (in mM) 150 NaCl, 5.4 KCl, 2.0 CaCl2, 2.0 MgCl2, 10 HEPES (pH 7.4), and 10 glucose. Patch electrodes (7-10 MΩ) were filled with (in mM) 120 CsCl, 10 BAPTA, 10 HEPES (pH 7.4), 4 MgCl2, and 2 ATP-Na2. After high resistance seal and break in (>1GΩ), whole cell voltage or current was recorded using an Axopatch 700B amplifier (Molecular Devices). Signals were low pass filtered (5 kHz) and digitized (5–40 kHz) using PClamp 10 software. For voltage clamp recordings, neurons were held at −60 mV. To record miniature AMPA excitatory post synaptic currents (EPSC); AP5, strychnine, bicuculline, and TTX were added to the external solution.

For LTD experiments, acute slices were prepared from hippocampus of wild-type, Flr or Flr injected animals, as previously described [25, 27]. Injections were performed at P17-P18 with HSV virus coding for SpCas9n and FL1 (sgRNAs). Recordings were performed 2-3 days later. Animals were anesthetized with isoflurane and decapitated. Brains were quickly removed and chilled in ice-cold dissection buffer. Transverse dorsal hippocampal 350-400 micron sections were cut using a VT-1000S vibratome (Leica, Germany) in ice cold carbogenated sucrose solution containing (in mM) 240 Sucrose, 2.5 KCl, 1 CaCl2, 5 MgSO4, 26 Na2HCO3, 1NaH2PO3, 11 Glucose, transferred to a chamber containing carbogenated artificial cerebrospinal fluid (ACSF) for 30 min at 32°C then maintained at RT (22-24°C) for a minimum of 1 hr prior to recordings. Electrodes were pulled to 3-5MΩ-tip resistance using a P-97 puller (Sutter Instruments, CA). Stimulating electrodes were fabricated tungsten bipolar electrodes (WPI) driven by ISO-STIM-01D (NPI Electronic Gmb). For recordings, slices were submerged and perfused (3 ml/min) in a carbogen-saturated ACSF at room temperature. Neurons positive for mRuby2 were selected for recording in the stratum radiatum of CA1. LTD was induced by paired-pulse low frequency stimulation (PP-LFS) consisting of 900 paired-pulses delivered at 1 Hz. Signals were amplified with a MultiClamp 700B, digitized with a Digidata 1440A, filtered at 2 kHz, sampled at 10 kHz and analyzed using pClamp 10 (Mol. Devices Corp., CA).

### Calcium events

Wild-type and Flailer primary neuronal cultures were infected at 3 DIV with AAV coding for GCaMP6s (Addgene #51086) and FL1 or control AAVs. 4-8 days after infection cells were placed into a temperature and CO2 controlled chamber (Tokai-Hit) over a Nikon TE-2000 epifluorescent microscope with a LED illumination system and a high speed Zyla camera (Andor, Ireland). Images were acquired every 30 ms for 10 minutes. Regions of interest were selected, and changes of fluorescence overtime calculated. Fluorescence peaks in 10 minutes were counted to calculate frequency of calcium events.

### Western blot analysis

Whole homogenate fractions were prepared from infected cultured neurons or brain tissue from wild-type, FL or FL-FL1 cells. Protein concentrations were determined using BCA (Pierce, IL). Proteins were separated on Any-KD PAGE gels (Biorad, USA) and transferred to PVDF membranes (Millipore, USA). Membranes were blocked and incubated overnight at 4C with primary antibodies. After rinsing, the membranes were incubated with secondary antibodies for 30 min at room temperature, rinsed and developed using chemiluminescence (Cell Signaling Technology, USA). Primary antibodies used: anti-MyoVa (LF-18; Sigma), anti-N-Cadherin (SC-7939; SantaCruz Biotech), Gnb5 (ab72406; Abcam). For detection HRP-conjugated secondary anti-bodies were used (Cell Signaling Technology, USA).

### RT-qPCR

Total RNA was extracted using Direct-zol RNA Miniprep Kits (Zymo Research, USA), according to manufacturer’s protocols. 400ng of total RNA was used for reverse transcription using LunaScript™ RT SuperMix Kit (NEB, USA). qPCR was performed using Forget-Me-Not™ EvaGreen® qPCR Master Mix (Biotium, USA). Data are presented as relative mRNA levels of the gene of interest normalized to GAPDH mRNA levels. Primers used for quantification: gnb5 sense 5’ GGGAACAAAGTCCTGTGCAT 3’, antisense 5’ GCGTGCTCCTTGTTCGTAGT 3’; myo5a sense 5’ TCCAGAAGCGTGTCACAGAG 3’, antisense 5’ CTTCTTCCTTTGCCTTGCTG 3’; flailer sense 5’ CAAAGGCCACGGGAACAAAG 3’, antisense 5’ CCAGAGGCACCTTCTTCTCA 3’; gapdh: sense 5’ ATGGTGAAGGTCGGTGTGAA 3’, antisense 5’ CATTCTCGGCCTTGACTGTG 3’.

### Stereotactic injection into the mouse brain

For stereotactic injections into the ventral hippocampus, P28-P30 wild-type and FL mice were deeply anesthetized by isoflurane administration delivered at 1-5%. Craniotomy was performed according to approved procedures. 700 nl of AAV PHP.eB mixture of FL1 (sgRNA (2.5×10^13^ vg/ml) + Cas9n (7×10^13^ vg/ml)) or tdTomato alone (3,5×10^13^ vg/ml) were injected at 100nl/min into the ventral hippocampus (anterior/posterior: −3.52; mediolateral: 2.65; dorsal/ventral: −3). Animals were sutured and analgesia was given (Buprenex, 1 mg/kg, i.p.). Animals were allowed to fully recover, and 3-4 weeks later behavioral experiments conducted. For electrophysiology recording we made use of HSV viral particles coding for sgRNA+Cas9n+mRuby3 (1×10^9^ infectious units/ml) or mRuby3 alone as control (1×10^9^ infectious units/ml). 300nl of HSV at 100nl/min was injected into the ventral hippocampus at P20. Analgesia was given (Buprenex, 1 mg/kg, i.p.) after surgery. 48-72 hours later acute brain slices were prepared.

### Intraventricular injections

Newborn wild-type and FL animals were anesthetized by hypothermia on an aluminum plate over ice maintaining the temperature above 1°C to avoid frostbite. 1μl of AAV PHP.eB mixture of FL1 (sgRNA (2.5×10^13^ vg/ml) + Cas9n (7×10^13^ vg/ml)) or tdTomato alone (3,5×10^13^ vg/ml) were injected into the cerebral ventricles bilaterally. Injection was performed using a 10 μl Hamilton syringe with a 32G beveled needle. Injection site was located half distance between bregma and lambda, 1 mm lateral to each side, and 3 mm deep. After injection animals were placed on a warming pad until they recovered color and regained movement to be later returned to their home cage.

### Brain sectioning and mounting

After behaviors to assess brain infection, animals were deeply anesthetized, and half of the brain was extracted and submerged to fix for a minimum of 24 hours in PBS CaMg + 4% PFA + 4% Sucrose into 30 mL flasks. After fixation, a Leica VT1000s vibratome was used to cut 100 μm coronal sections. Slices were kept in PBS and mounted using Fluoromont G (EMS, Hatfield, PA) to preserve the fluorescence signal. Brain images were captured with a Nikon Eclipse TE2000 epifluorescence microscope or a Nikon C2+ Confocal (Nikon, USA).

### Behavioral testing

For all behavioral testing, P28-P30 male and female wild-type and FLR animals were injected as previously described with AAVs coding for tdTomato alone (control) or together with FL1 (Cas9n +sgRNA). 8 weeks after infection animals were used for behavioral testing as has been previously described [27].

### Grooming behavior

Animals were individually placed into a new cage and allowed to habituate for 20 minutes. After habituation, animals were filmed for 1 hour between 19:00 and 21:00 hours with a red light (2 lux) in a Behavior Spectrometer testing chamber (Behavioral Instruments). Grooming was quantified automatically and corroborated manually using the Viewer software. The total time spent grooming in a 1-hour interval was determined. Grooming included all sequences of face-wiping, scratching/rubbing of head and ears, and full-body grooming.

### Elevated plus maze test

For the elevated plus maze, open arms were illuminated by 60 lux light while the dark (closed) arms were lit by 10-20 lux. Before testing, animals were left to habituate to 10-20 lux light for 1 hour. For testing, animals were placed in the center of the maze and allowed to explore the maze freely for 5 minutes, all movements were recorded. Analysis was performed using the automated tracking software Noldus Ethovision. Anxiety-like behavior was determined by the time spent in the open arm during the 5 minute interval.

### Light and dark test

Mice were habituated to 20-40 lux light 1 hour prior to testing. Testing was conducted in a 2-chamber apparatus (Med Associates) where one side was left completely dark (dark chamber) and the other illuminated at 1000 lux (light chamber) with an overhead lamp. Mice were placed in the dark chamber and a connecting door between dark and light chambers opened. Mice were left to freely explore for 5 minutes. The latency to emerge to the light chamber, the number of crosses and the total time spent in the light chamber were automatically scored by Noldus Ethovision software. Anxiety-like behavior was determined by the time spent in the light chamber, and the latency to enter it in a 5-minute interval.

### Fear conditioning

Animals were habituated to the experimenter and behavioral room 2 days previous to testing. Animals were taken to behavioral room and placed into an isolated fear conditioning chamber (Med Associates Inc.), left to habituate for 8 minutes for 2 consecutive days, and returned to their home cage. At Day 3 (training day), animals were placed on the same chamber and let explore freely for 2 minutes followed by a 2 second 0.6 mA mild shock. After shock, animals were left to explore for 3 more minutes and returned to their home cage. At day 4 (testing day), mice were placed in the conditioning context and freezing to the context provided by the box alone was assessed over 5 min. Freezing behavior was recorded, and analysis was performed using FreezeFrame (Actimetrics, USA).

### Data analysis

All values are presented as mean ± standard error (SE). Number of experiments, animals or cells are indicated in each figure legend. Data normality was checked using the Shapiro-Wilk test. Statistical analyzes were performed using one-way ANOVA followed by Bonferroni post-hoc test. Values of p<0.05 are considered statistically significant. *p<0.05, **p<0.01, ***p<0.001. All statistical analyzes were performed using Graphpad Prism.

## Authors’ contributions

FJB design the work, acquired, and analyzed data, wrote the manuscript. SP performed behavioral experiments. HH acquired data and analysis. JPZ performed electrophysiological recordings. HS analyzed data. MH and LS helped with experimental design and acquired data. BD and VG helped with AAV experiments and provided materials. FZ design the study. MCP conceptualized and designed the study.

## Declarations

All animal protocols were approved by MIT Committee on Animal Care (CAC), and Ethical and Bio-security Committees of Universidad Andres Bello.

All data and materials are available upon request.

## Acknowledgements

We thank Lorena Varela-Nallar and Gloria Arriagada for critical reading of the manuscript and helpful feedback.

The authors declare that they have no competing interests

---

**Supplementary Figure 1.**
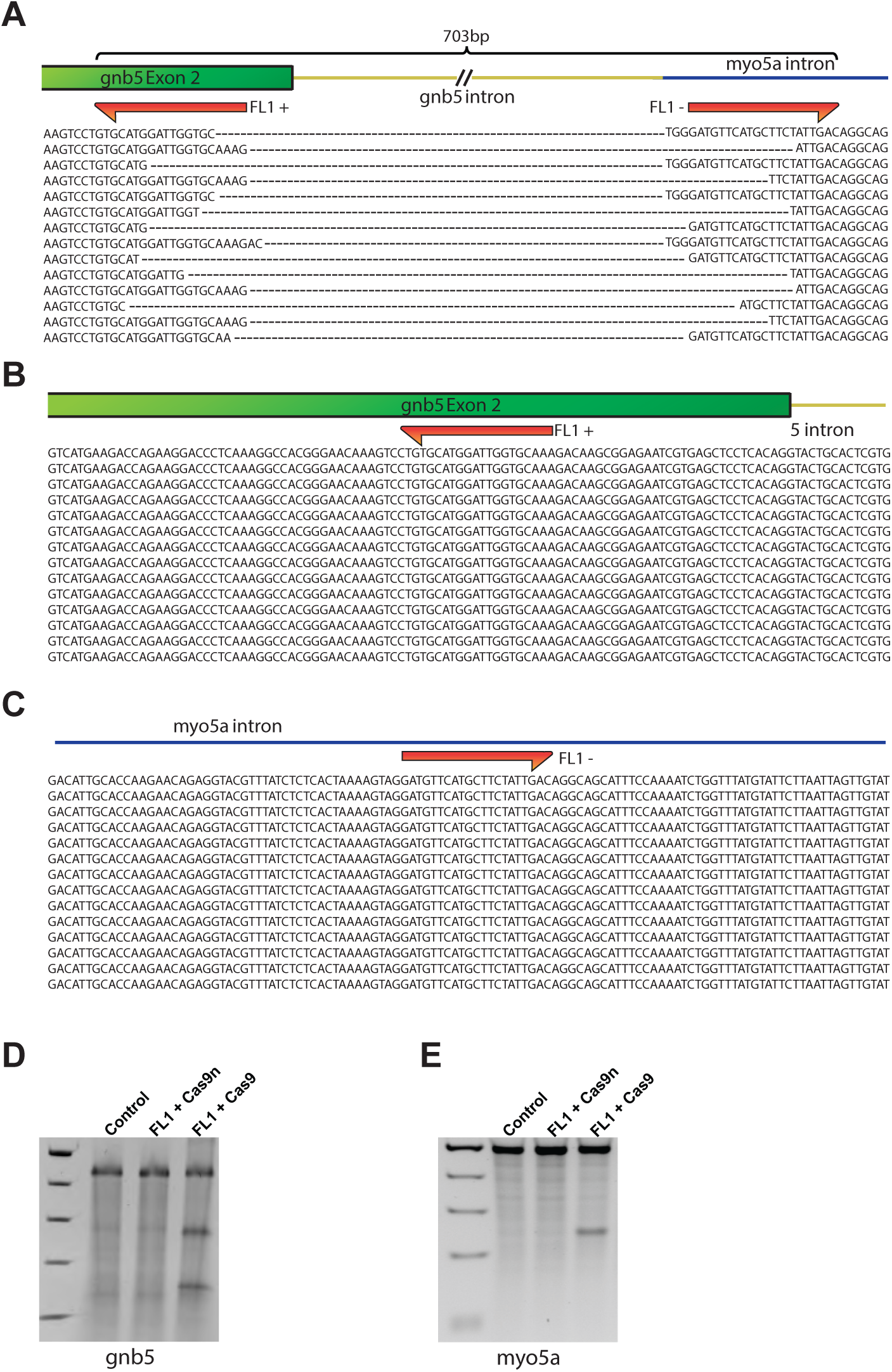
FL1 DN-CRISPRs successfully target the flailer genomic locus without targeting gnb5 or myo5a. (A) Representative sequences of PCR products sequenced to determine changes in genomic sequence at the flailer locus after FL1 infection. (B-C) Representative sequences of gnb5 (B) and myo5a (C) genomic locus showing no alteration in the sequences were FL1 DN-CRISPRs target the endogenous genes. (D-E) SURVEYOR assay to analyze gene editing of gnb5 (D) or myo5a (E) shows no edition at the locus of endogenous genes when Cas9n is used. Note that co-infection of FL1 and Cas9 produces double strand breaks and edition of gnb5 and myo5a.

**Supplementary Figure 2.**
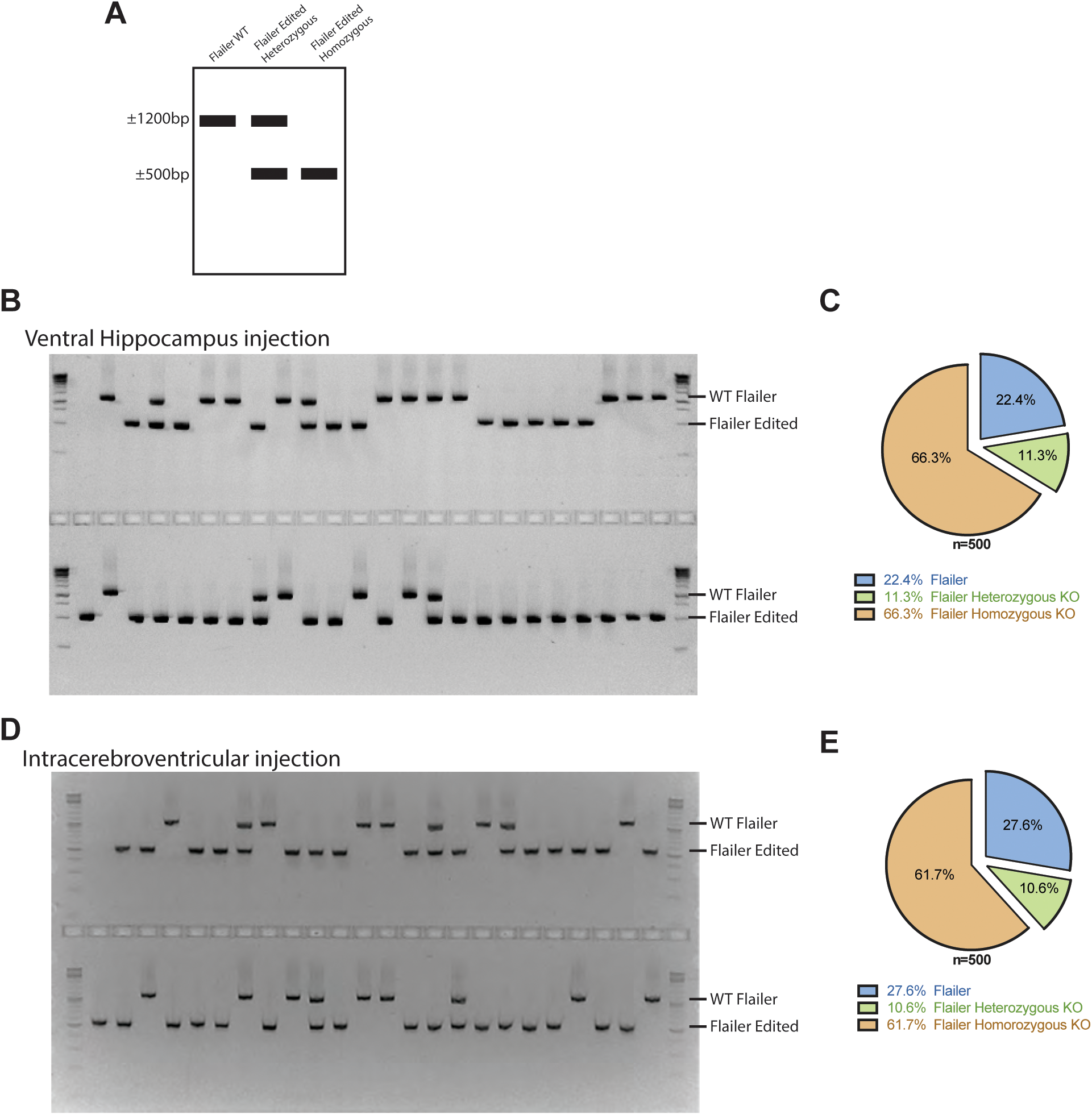
Single cell PCR to determine gene edition in FL1icv injected Flailer animals. (A) Scheme depicting the expected PCR products of FL animals (∼1200bp), FL edited heterozygous (∼1200bp + ∼500bp), and FL edited homozygous (∼500bp). (B) Representative gel of PCR products from single cell FL1 ventral hippocampus infected tissue. (C) Quanti cation of the number of FL, FL heterozygous, and FL homozygous products. A total number of 500 cells were analyzed from 10 FL1icv animals. (D) Representative gel of PCR products from single cell FL1 cortical infected tissue by intracerebroventricular injections. (E) Quanti cation of the number of FL, FL heterozygous, and FL homozygous products. A total number of 500 cells were analyzed from 10 FL1icv animals.

